# SpeciFlex: A Protocol for Mining Binding Site Conformational Selectivity in Structure-Based Inhibitor Discovery

**DOI:** 10.1101/304931

**Authors:** Matthew E. Tonero, Leslie A. Kuhn

**Author notes:** Author for correspondence: Department of Biochemistry & Molecular Biology and Computer Science & Engineering, Michigan State University, East Lansing, MI 48840 USA, Web: http://www.kuhnlab.bmb.msu.edu.

## Abstract

Selectivity for a target site is challenging when the site is conserved in homologous proteins. A novel protocol is presented for attaining selectivity by taking advantage of conformational population differences between homologs. Conformational ensembles of the targeted protein and the homolog are compared to identify pockets significantly populated in the target, but rarely or never sampled in the homolog. SLIDE screening then identifies molecules that fit the unique pocket and also interact well with an adjacent substrate pocket. The SpeciFlex protocol, demonstrated for a pair of pyrophosphokinases, yields ligand candidates with good interactions in both the substrate and unique pockets of the target *Yersinia pestis* protein, while selecting against interactions with the homologous site in *Escherichia coli*.

## Introduction

### Conformational selection

Conformational change occurs, to a greater or lesser extent, when all proteins and ligands form complexes. The rigid lock-and-key-fit model, with the protein as the lock and the ligand as the key, has been shown inadequate to model the transition between most ligand-free and bound protein structures, leading to a new focus on modeling molecular flexibility upon complex formation [1–9]. The conformational selection hypothesis [10–11], arising from energy landscape concepts in protein folding theory [12–13], describes the native state of the protein as sampling an ensemble of conformations between the ligand-free (open) and ligand-bound (closed) states. A ligand can select from this array of conformations and bias the equilibrium toward the catalytically competent or inhibitor-bound conformation. Several proteins have been shown to sample the conformations required for ligand binding as part of their native ensemble, including staphylococcal nuclease [14–15], calbindin [16], adenylate kinase [17–18], and calmodulin [19]. A central concept here is that the energy required to shift the conformation of either the protein or ligand into the bound state becomes a penalty that decreases their affinity of interaction. Therefore, requiring an energetically expensive conformational change is unlikely.

### Conformational differences as the basis for ligand selectivity

While the amino acids within binding sites tend to be highly conserved among structurally homologous proteins employing the same substrates and carrying out the same chemical reactions, near-closed conformations of these enzymes can differ. This is due to amino acid sequence variation that occurs near the binding site, particularly in binding-site loop side chains that face away from the site, which influence conformational sampling between the closed and open states. Furthermore, there is decreased evolutionary selection on these non-catalytic conformations. Statistical mechanics and the Boltzmann distribution tell us that the relative prevalence of conformations reflects their relative favorability in energy. Thus, for a targeted conformation to be useful as a template for discovering selective inhibitors, it is not necessary that the conformation be absolutely unique in the targeted protein relative to a homolog, but that it be well-sampled (low in energy) in the targeted protein and rarely sampled (much less favorable in energy) in the homolog.

A well-known example of selectivity gained by targeting a unique conformation is the cancer drug, Gleevec (imatinib), which was found by X-ray crystallography to bind to a unique, inactive kinase conformation, locking the activation loop. Closely related Src kinases cannot adopt this inactive conformation, allowing Gleevec to attain high selectivity for its target, Abl kinase [20]. While this was a fortuitous discovery for Gleevec, the method presented here is designed to identify pockets conferring conformational selectivity, *a priori*. The search for ligands that selectively bind one protein relative to its homologs can then be directed towards the unique conformation.

We show that such conformations can be used as templates for SLIDE screening to discover ligands that are target-selective. While other screening methods such as FlexE [21] have also employed multiple protein conformations (typically from independent crystal structures) as input to docking and screening, and molecular dynamics analysis has been used to identify conformationally invariant regions as targets for drug discovery [22,23], the idea here is take advantage of differences in conformational occupancy sampled by protein dynamics as a way of enhancing target selectivity.

### Biology of hydropterin pyrophosphokinases

This study focuses on the folate biosynthetic enzyme, 6-hydroxymethyl-7,8-dihydropterin pyrophosphokinase (HPPK), a target for antibiotic development. Mammals use active transport for taking up folate compounds from their diets, while most microorganisms synthesize folates. Thus, the folate biosynthetic pathway is one of the principal targets for developing antibiotics [24]. HPPK catalyzes the transformation of 6-hydroxymethyl dihydropterin to 6-hydroxymethyl dihydropterin pyrophosphate via the transfer of pyrophosphate from ATP. Here we focus on the identification of inhibitors specific to pathogenic *Yersinia pestis* HPPK, relative to the beneficial gastrointestinal bacterium, *Escherichia coli. Y pestis* is the causative agent of bubonic, pneumonic, and septicemic plague and one of the most virulent pathogens known [25]. Three recorded plague pandemics are estimated to have killed 200 million people [26]. The recent emergence of a multi-drug resistant strain of *Y pestis*, resistant to all first-line and several prophylactic antibiotics, is of particular concern [27].

Our study begins with the crystallographic closed, ternary complexes of *Y pestis* and *E coli* HPPK bound to AMPCPP (an ATP analog in which the oxygen joining AMP to pyrophosphate is replaced by a non-hydrolyzable, single-bonded carbon), 6-hydroxymethy-7,8-dihydropterin (the other natural substrate), and two Mg^2^+ ions, as deposited in Protein Data Bank [28] (PDB) entries 2QX0 and 1Q0N [29–30]. There are only two amino acid differences between *E coli* and *Y pestis* HPPK near the targeted dihydropterin binding pocket. One is the *E coli* Tyr53 →*Y pestis* Phe difference, in which the hydrophobic contact made by these two side chains is conserved, and the hydroxyl group of the Tyr residue interacts with bulk solvent. The other is the *E coli* Pro43 →*Y pestis* Lys difference at the base of a binding site loop, with the side chain directed away from the dihydropterin. The result is that the dihydropterin pocket is highly conserved in closed structure and chemistry between the two species, providing no static differences that can be employed to attain inhibitor specificity. This makes the *E coli* and *Y pestis* dihydropterin sites in HPPK an ideal model system for exploring the use of conformational sampling differences as the basis for discovering species-selective inhibitors.

### Experimental Methods

The SpeciFlex1 protocol is divided into two steps: conformational analysis to identify a uniquely available pocket near the *Y pestis* dihydropterin site (Step 1, summarized in Figure 1) and virtual screening to identify compounds that interact well with the unique pocket in *Y pestis* but cannot interact similarly well with *E coli* (Step 2, summarized in Figure 2). In this particular application, compounds that also mimic dihydropterin interactions with *Y pestis* HPPK are sought, because ligand candidates that share interactions with known inhibitors have an enhanced probability of inhibition [31]. However, the method is entirely general and could be applied to the neighborhood of any known ligand pocket, or in fact to any binding cleft relevant to regulating protein activity (potentially an allosteric site). Following are details of the methods, which are cross-referenced by using the same step labels in the flowcharts (Figures 1 and 2). The Results then show how these steps have been applied to HPPK to discover potential inhibitors that meet specific criteria for quality of interaction with the dihydropterin and unique pockets in *Y pestis*, while taking advantage of the differences in protein dynamics between species.

**Figure 1.**
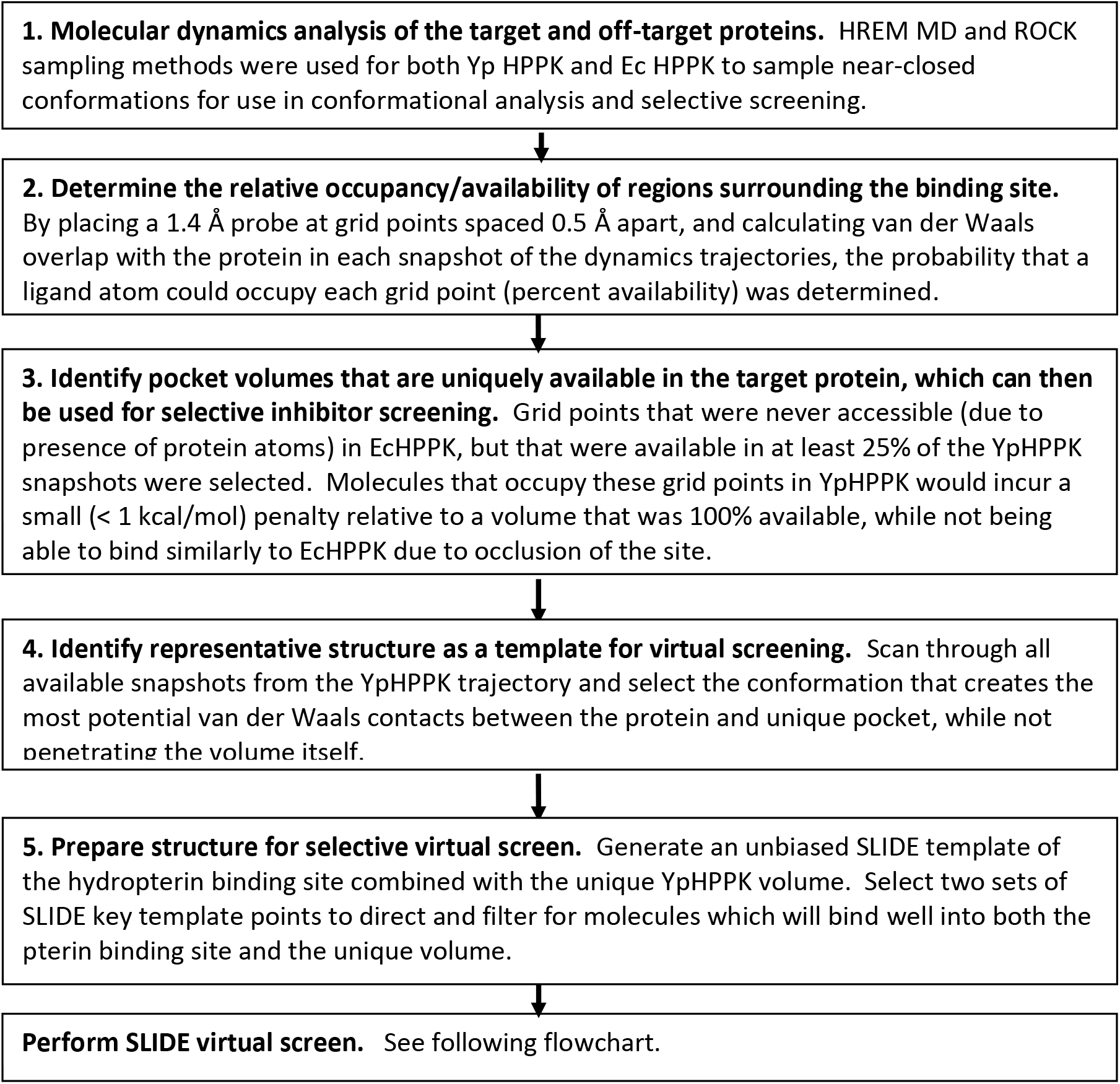
SpeciFlex Part 1: Analyzing molecular dynamics simulations to identify a pocket adjacent to the substrate site that is uniquely available in the targeted protein.

**Figure 2.**
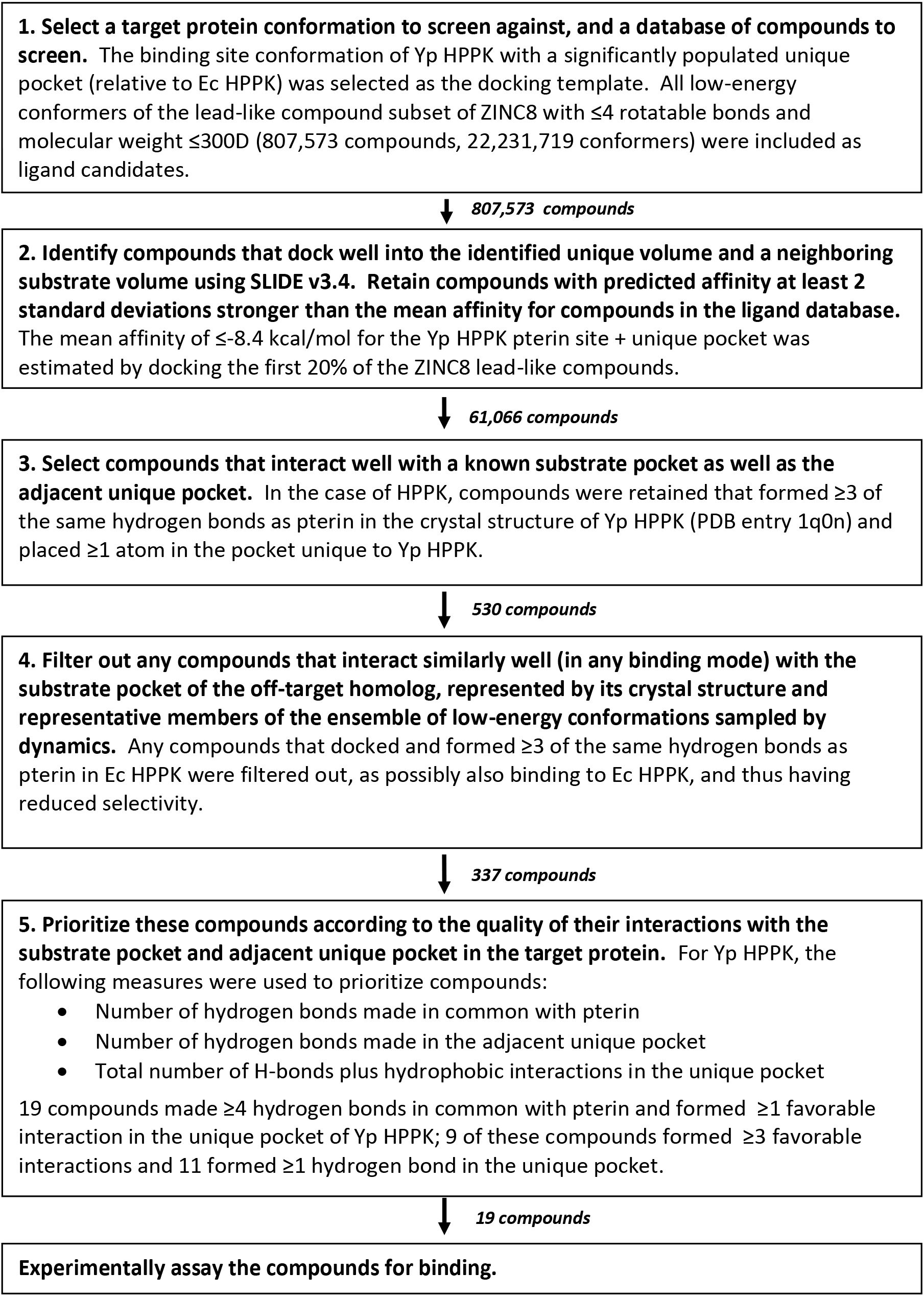
SpeciFlex Part 2: Identifying ligand candidates that are selective for the unique pocket and also interact well with the substrate site.

### Step 1-1. Molecular dynamics conformational analysis of the protein target and its homolog

The starting point for the SpeciFlex methodology (step 1-1, Figure 1) is the analysis of molecular dynamics trajectories that sample conformations near the substrate-bound, closed state of the binding site in the target protein as well as the off-target homolog. These trajectories can arise from molecular dynamics, elastic network models [32], or methods like ROCK that sample dynamics consistent with maintaining a set of non-covalent interactions, e.g., those present in the closed structure [33–34]. Given that any simulation method has sampling biases arising from the choice of potential energy function and parameterization of atoms and bonds, we chose to combine two simulation approaches: a fine-grained molecular dynamics simulation with explicit solvent, and a coarse-grained network model. By determining the consensus between the two dynamics trajectories in terms of volumes near the dihydropterin binding site available for ligand binding, the aim was to ensure that the identification of a pocket unique to the target protein remains robust with respect to variations in the way the dynamics are sampled.

#### Molecular dynamics analysis

Explicit-solvent molecular dynamics trajectories were provided for *Y pestis* and *E coli* HPPK by colleagues Li Su and Robert Cukier (Department of Chemistry, Michigan State University), starting from the closed, ternary complexes (PDB entry 1Q0N for *E coli* and PDB entry 2QX0 for *Y pestis*) from which dihydropterin had been removed and AMPCPP was replaced by the native ATP, modeled using the GROMOS9644 force field [35]. Using the Hamiltonian replica exchange method (HREM), replicas of the system that differed in potential energy terms were simulated concurrently, and system configurations were exchanged periodically. Three terms in the potential energy function had Lennard-Jones and electrostatic nonbonded interactions scaled differently between the replicas to enhance conformational sampling. These terms represented the potential energy of interactions (i) between atoms in loops 2 and 3, which close over the dihydropterin and ATP sites in HPPK, (ii) between a loop atom and an atom in the rest of the protein (largely a scaffold-like beta sheet structure), and (iii) between the atoms in the rest of the protein [35]. A small number of replicas were found to be sufficient to enlarge the sample space substantially, on the basis of root-mean-square fluctuation measures. The result is enhanced movement through configuration space while maintaining Boltzmann sampling. Snapshots taken every 1 picosecond from nanoseconds 3 through 5 of the *Y pestis* and *E coli* HPPK simulations (post-equilibration) were used for subsequent conformational analysis.

The next goal was to identify shared features in the conformational ensembles identified by HREM molecular dynamics and Rigidity Optimized Conformational Kinetics (ROCK) simulations [33–34]. ROCK samples motions consistent with maintaining a set of non-covalent interactions in the protein, in this case the hydrogen bonds, salt bridges, and hydrophobic interactions in the ligand-free closed state of HPPK. In the new version (2.6) of ROCK, only stereochemically favorable side-chain and main-chain dihedral angle changes are sampled in low-energy motions of underconstrained regions. Non-covalent interactions for ROCK were identified using ProFlex (v5.1), which uses the graph theory approach FIRST [36–37] to determine the network of covalent bonds, hydrogen bonds, salt bridges, and hydrophobic interactions and identify regions of the protein that are under-constrained and therefore flexible.

#### Details of ProFlex analysis

The A chains of the two PDB entries (1Q0N for E coli HPPK, and 2QX0 for *Y pestis*) were prepared for ProFlex analysis by removing all ligands and water molecules and using the most highly occupied state in the case of atoms with multiple occupancies. The magnesium ions were treated as charged hydrogen bond donors. All hydrogen bonds, salt bridges and hydrophobic tethers were kept, subject to a hydrogen-bond energy threshold of −1.967 kcal/mol for *E coli* and −1.885 kcal/mol for *Y pestis* HPPK. At these energy levels, the rigid core and flexible regions in HPPK matched experimentally observed data (the NMR ensemble and crystal structures determined along the catalytic cycle of HPPK), as well as maintaining an average of 2.4 covalent + non-covalent interactions per atom, which is characteristic of the bond networks of the native states of proteins in general [38].

#### Details of ROCK analysis

ROCK trajectories were defined using the following parameter settings, most of which govern the efficiency of sampling: ring cluster rotation ratio (percentage of bonds sampled in a given step): 10%; ring-cluster rotation limit (maximum rotation of any bond per step): 5.0 degrees; Ramachandran region (ProCheck region of main-chain dihedral space to be sampled; [39]): generous; van der Waals coefficient: 0.6 (van der Waals overlaps are only detected when two atoms are within the sum of their radii multiplied by this coefficient; however, only collision-free conformations are used at the end of the ROCK trajectory, based on CHARMM energy minimization [40]); output step: 8000 conformations sampled, with every 20^th^ structure output, for a total of 400 structures. These parameters resulted in fine-grained sampling of conformations near the closed state. The energy minimization of ROCK conformers employed CHARMM v. 31b1 with the CHARMM22 force field and generalized Born solvation for 300 steps to remove any unfavorable steric or electrostatic interactions.

#### Superposition of HPPK conformers

For comparing HREM snapshots and ROCK conformers of *E coli* and *Y pestis* HPPK, rigid regions defined by HREM analysis [35] were least-squares fit to overlay the molecules, as some tumbling or shifting of molecules can occur during dynamics simulation. The main-chain atoms of residues 1-9, 15-42, 54-79, 93-100, 125-132, and 143-158 (PDB entry 1Q0N), which form a relatively rigid core in *E coli* HPPK, were least-squares superimposed onto those of residues 2-10, 16-43, 55-80, 94-101, 126-133, and 144-159 of *Y pestis* HPPK (PDB entry 2QX0).

### Step 1-2. Determine the relative occupancy/availability of volumes surrounding the binding site

The HREM and ROCK trajectories of *E coli* and *Y pestis* HPPK were considered individually (as four cases), with a 0.5 Å grid placed over the superimposed trajectories. For each grid point, the percentage of the trajectory that a probe ligand atom of 1.4Å radius could occupy grid points in or near the dihydropterin site (without interpenetrating the protein) was calculated.

The neighborhood of the dihydropterin site evaluated was within 10 Å of the position of dihydropterin in the closed crystal structure in the search for volumes available in *Y pestis* but not *E coli* HPPK. Isocontouring using OpenEye VIDA software was applied to the grid values to define surfaces encompassing volumes representing 0%, 25%, 50%, 75%, and 100% availability for ligand atoms from each trajectory. The trajectories were then combined for *Y pestis* and separately for *E coli HPPK*, to identify volumes that were accessible 0%, 25%, 50%, 75% or 100% or the time in both the ROCK and HREM trajectories for the target and homolog, respectively. This presents a more stringent criterion for identifying an available (or disallowed) volume, since it had to be present in both dynamics trajectories.

### Step 1-3. Identify pocket volumes that are uniquely available in the target protein, which can then be used for selective inhibitor screening

The ideal case is when a volume near the known substrate (e.g., dihydropterin) site is available 100% of the time in the target protein but never available (0%) in the off-target homolog. Given the proteins’ homology, however, it may be difficult to find volumes of meaningful size that meet these stringent criteria. In the next simplest case, a pocket is available 0% of the time in the offtarget homolog, and available X% of the time in the target. In this case, the selectivity is absolute, but the energy to reach that conformation in the target must be small enough to not subtract much from the binding affinity, and the inhibitor must absolutely complement that volume and not bind to the off-target homolog in another favorable way. This case is considered in depth in the Results section.

In this section, the general case is considered in which a pocket is somewhat accessible in both the target and its homolog. The relative pocket availability across the trajectory, reflecting underlying conformational populations, influences the free energy of binding to that pocket. The difference in binding affinity of the ligand for the target relative to the homolog will be the result of differences in protein-inhibitor interactions based on different functional groups being exposed in the open volume (which is measured by the affinity prediction scoring function), as well as the relative probability of the pocket volume being accessible in the target versus the homolog. This is proportional to the log of the relative populations, as follows:

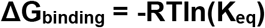

where

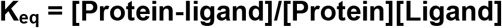

If the population of the binding site conformation required to bind a specific ligand (the target conformation) is considered in relation to all conformations of the protein, then the prevalence of this target conformation (its “concentration” in solution) can be determined. The concentration of the target conformation can then be compared for different homologs of a protein, to determine the difference in free energy needed to reach the target conformation. For two homologs, A and B, the contribution to ΔG_binding_ from the difference in concentration of the target conformation can be calculated as:

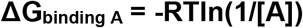

and

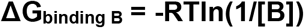

where [A] is the concentration of the target conformation in homolog A, and [B] is its concentration in homolog B. So, the difference between binding free energies for the two homologs is:

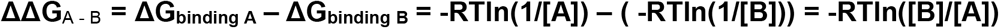

Given room temperature (T = 293K) and the value of the ideal gas constant (R=1.986 cal/mol*K):

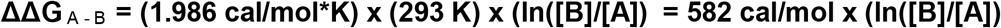

Thus, a 1.34 kcal/mol difference in ΔG_binding_ occurs for every ten-fold difference in population between homologs A and B. This only considers the availability of a pocket, and not other potential factors affecting binding affinity, such as differences in binding site interactions and solvation, which can be evaluated by the affinity scoring function. The binding free energy cost of attaining conformations containing the desired pocket, as a function of prevalence of those conformations, is provided in Table 1. When a pocket is rarely available (e.g., only 0.02% of the time), the penalty in ΔG for achieving such conformations becomes quite significant, about 5 kcal/mol. However, for a targeted conformation that is found in 20% of the population, the penalty in change in free energy to reach that conformation is less than 1 kcal/mol. Thus, it is reasonable to aim for a pocket that is present in 20-100% of the protein’s conformations and is never or very rarely sampled (e.g., 0.02%) in the off-target homolog. This unique pocket can then targeted for inhibition along with a neighboring substrate pocket.

**Table 1.** The cost in binding free energy of attaining a conformation in which the targeted unique pocket is available. If the pocket is always available in the target protein, A, there is no cost, and if the pocket is only represented in 0.01% of the dynamics trajectory, the cost is substantial (5.36 kcal/mol). These pocket availability values can be used to determine the energetic difference of binding to protein A relative to protein B when protein A contains the pocket X% of the time and protein B contains the pocket Y% of the time. For example, if the pocket is 60% available in protein A and only 10% available in B, the free energy difference corresponding to pocket availability would be 0.30 kcal/mol - 1.34 kcal/mol, making binding to the pocket in protein A 1.04 kcal/mol more favorable. This calculation does not take other enthalpy or entropy factors into consideration (which are considered by the scoring function used for docking), and is based solely on pocket availability.

| Availability of Pocket Volume (% of trajectory) | Cost in $\Delta G_{\text{conf}}$ (kcal/mol) | Availability of Pocket Volume (% of trajectory) | Cost in $\Delta G_{\text{conf}}$ (kcal/mol) |
| --- | --- | --- | --- |
| 100 | 0 | 0.9 | 2.74 |
| 90 | 0.06 | 0.8 | 2.81 |
| 80 | 0.13 | 0.7 | 2.89 |
| 70 | 0.21 | 0.6 | 2.98 |
| 60 | 0.30 | 0.5 | 3.08 |
| 50 | 0.40 | 0.4 | 3.21 |
| 40 | 0.53 | 0.3 | 3.38 |
| 30 | 0.70 | 0.2 | 3.62 |
| 20 | 0.94 | 0.1 | 4.02 |
| 10 | 1.34 | 0.09 | 4.08 |
| 9 | 1.40 | 0.08 | 4.15 |
| 8 | 1.47 | 0.07 | 4.23 |
| 7 | 1.55 | 0.06 | 4.32 |
| 6 | 1.64 | 0.05 | 4.42 |
| 5 | 1.74 | 0.04 | 4.55 |
| 4 | 1.87 | 0.03 | 4.72 |
| 3 | 2.04 | 0.02 | 4.96 |
| 2 | 2.28 | 0.01 | 5.36 |
| 1 | 2.68 | 0.009 | 5.42 |

### Step 1-4. Identify a representative structure as the template for virtual screening

All 2400 HREM and energy-minimized ROCK snapshots of the target protein were considered in the context of the unique volume and adjacent substrate site defined in the previous step. By cycling through the snapshots one by one using a computational script, the conformation was chosen that yielded no steric overlaps and provided the maximum number of favorable van der Waals contacts with the substrate and unique pocket volumes. This conformation was used as the target for selective virtual screening.

### Step 2-1. Preparing the database of ligand candidates for virtual screening

All low-energy conformers of the lead-like compound [41] subset of ZINC8 [42]; http://zinc.docking.org), filtered to have ≤4 rotatable bonds and molecular weight ≤300D, were included as ligand candidates. Compounds with no more than four rotatable bonds were the focus, because highly flexible molecules pay a large entropic cost upon binding, require disproportionate screening time to represent their entire conformational space, and their flexible groups often make relatively few good interactions. In selecting compounds with molecular weight of 300D or less, the rationale was that compounds about 50% larger than the native substrate, 6-hydroxymethyl-7,8-dihydropterin, with a molecular weight of 195D, could substantially fill the pterin binding site and also make significant interactions in the unique pocket, while remaining amenable to optimization by addition of chemical side chains that enhance protein interactions.

Small compounds tend to be better starting points for chemical optimization, since it is often easier to add groups that optimize interactions rather than remove suboptimal parts of larger compounds. Version 2.3.2 of Omega (OpenEye Scientific Software, Santa Fe, NM) was used with default settings to sample all 22,231,719 low-energy conformers of the 807,573 commercially available compounds meeting the above criteria (an average of 28 conformers per compound).

### Step 2-2. Identify compounds that dock well into the identified unique volume and a neighboring substrate volume using SLIDE

The first step in screening by SLIDE version 3.4 [43–44] (http://www.kuhnlab.bmb.msu.edu/software/slide/index.html) involves automated definition of a docking template. This represents positions in the ligand binding cavity where ligand atoms can make favorable hydrogen-bond, salt-bridge, or hydrophobic interactions with atoms in the target protein. Only information from the protein was used in determining the positions of template points. In the case of the dihydropterin pocket, template points were placed within the volume falling within 3 Å of dihydropterin bound to *Y pestis* HPPK (PDB entry 2QX0) to ensure the pterin pocket was fully sampled. The ligand was used only to define the volume of the pocket, and was not used for positioning the template points. The unique volume identified next to the pterin pocket (Figure 1, step 3) was also populated with template points automatically. Candidate ligands are permitted to interact with the protein well outside the template region in SLIDE, and the quality of interaction for each docked molecule is measured by an affinity scoring function that considers interactions between all protein and ligand atoms. To ensure that each docked ligand interacts with the dihydropterin pocket (known to provide good protein interactions) and also the unique volume (favoring compounds selective for *Y pestis*), the key point option in SLIDE was used. SLIDE version 3.4 allows the user to specify up to two sets of key points (user-defined subsets of the entire set of template points); any docking must match at least one key point in each set. The first set was defined as template points placed positioned deep in the dihydropterin pocket, relative to its C=C bond. The second set comprised all template points within the *Y pestis* HPPK unique volume defined by consensus of the HREM MD and ROCK dynamics.

Given dockings that were free of steric overlap and interacted with both the pterin and unique pockets, the overall quality of docking was evaluated by atomistic semi-empirical protein-ligand scoring. SLIDE versions 3.0 and above include two-tiered scoring to identify the optimal binding mode of each molecule in the first step using OrientScore, and then predict the affinity of the docked molecule in a second step using AffiScore [44]. SLIDE 3 docking using OrientScore followed by one second of energy minimization with Szybki (OpenEye Scientific Software) has been found to perform as well as GOLD and Surflex (the closest competitors), and somewhat better than GLIDE, FlexX, DOCK, QXP, and FRED, in accurately re-docking ligands in 100 protein complexes; 51% of the ligands were docked to within 1 Å RMSD and 54% to within 1.5 Å RMSD [44–45]. The AffiScore scoring function used to predict ΔG_binding_ by SLIDE had a linear correlation coefficient of 0.65 with the experimentally determined binding affinity values for 273 complexes, similar to the best method tested (XScore; [46]), which had a linear correlation coefficient of 0.68 [44]. In discovery applications, SLIDE identified seven new classes of low to mid-micromolar inhibitors for the ATP binding site in asparaginyl-tRNA sythetase from the human parasite *Brugia malayi*, with a hit rate of 15% [50]; 7 out of 45 compounds assayed were confirmed as inhibitors. Flexibility modeling in this system also helped explain why a series of long side-chain analogs of the top-scoring inhibitor proved to be selective for the *Brugia* target relative to its human homolog, despite the absence of amino acid differences within the binding pocket. In a thrombin inhibitor discovery collaboration, 5 out of 26 (19%) of the top-scoring soluble SLIDE compounds were found to inhibit human thrombin, with IC_50_ values in the mid-nanomolar to mid-micromolar range [31]. Thus, we anticipate a hit rate of 15-19% in SLIDE applications, though that may depend significantly on the enzyme, ligand database, and scoring approach.

For retaining the best candidates from the 22 million ZINC8 lead-like conformers, an affinity threshold was set following screening the first one-fifth of the database. Only those compounds with an AffiScore at least two standard deviations better than the mean score (≤-8.4 kcal/mol) were kept, upon docking to *Y pestis* HPPK.

### Step 2-3. Select compounds that interact well with both the known substrate pocket and the adjacent unique pocket

Given that a large number of compounds (61,066) met all the above criteria (Figure 2), and current affinity prediction scoring functions have mean errors of ~2 kcal/mol, additional criteria were considered for selecting inhibitors. Previous results [31] indicate that high-scoring molecules identified by SLIDE that were subsequently proven to inhibit thrombin tended to share many interactions with known inhibitors. Other groups have explored the same approach in terms of filtering dockings according to pharmacophore constraints [51,52]. Conserved hydrogen-bond and salt bridge interactions provided the most discrimination. Thus, for HPPK, compounds were retained that formed ≥3 of the same hydrogen bonds as dihydropterin in the crystal structure of *Y pestis* HPPK (PDB entry 2QX0) and placed ≥1 atom in the unique pocket.

### Step 2-4. Filter out compounds that interact similarly well (in any binding mode) with the substrate pocket of the off-target homolog

Compounds that bound well to *Y pestis* HPPK using the above criteria are virtually guaranteed to not bind in the same orientation to any conformation of *E coli* HPPK. This is because the unique pocket volume was absent in the *E coli* closed crystal structure and all its dynamics conformations. However, a compound could conceivably bind to *E coli* in a different conformation or orientation and still make good interactions with the dihydropterin pocket, diminishing its selectivity for *Y pestis* HPPK. Thus, any compounds that docked and formed ≥3 of the same hydrogen bonds as pterin in *E coli* HPPK (PDB entry 1Q0N) were filtered out.

### Step 2-5. Prioritize these compounds according to the quality of their interactions with the substrate pocket and adjacent unique pocket in the target protein

Given the remaining limited number of compounds (337, in the case of the HPPK screen), all of these compounds could be assayed for inhibition and other biological activities. Alternatively, they can be prioritized further by other protein-ligand scoring functions (e.g., DrugScore, EON, or GOLD, as was performed in [31]). This choice is typically based on the software available to the user and any background knowledge of which scoring function(s) work best for that protein family. Here, based on our prior experience, they were prioritized based on the: i) number of hydrogen bonds in common with a known ligand, dihydropterin [31], ii) number of hydrogen bonds made in the adjacent unique pocket, and iii) total number of hydrogen bonds plus hydrophobic interactions in the unique pocket. These proved useful for focusing in on compounds exhibiting protein complementarity typical of what a structural biologist would expect to see in a protein-inhibitor complex.

## Results and Discussion

### Molecular dynamics analysis

Sampling achieved by HREM MD analysis for *E coli* HPPK and *Y pestis* HPPK (Figure 3) indicated extensive sampling of loop conformations surrounding the dihydropterin binding site. For ready visualization, the 2000 structural snapshots taken 1 ps apart during the equilibrated 2 ns dynamics trajectory were clustered according to main-chain conformation, and representative conformations are shown for each cluster (Figure 3). ROCK dynamics sampled closely around the closed crystal structure of HPPK with dihydropterin removed. The average pair-wise RMSD value for main-chain atoms in loops 2 and 3 across all 400 ROCK conformations (representing 8000 accepted conformational steps) was 1.39 Å for *Y pestis* HPPK and 0.73 Å for *E coli* HPPK.

**Figure 3.**
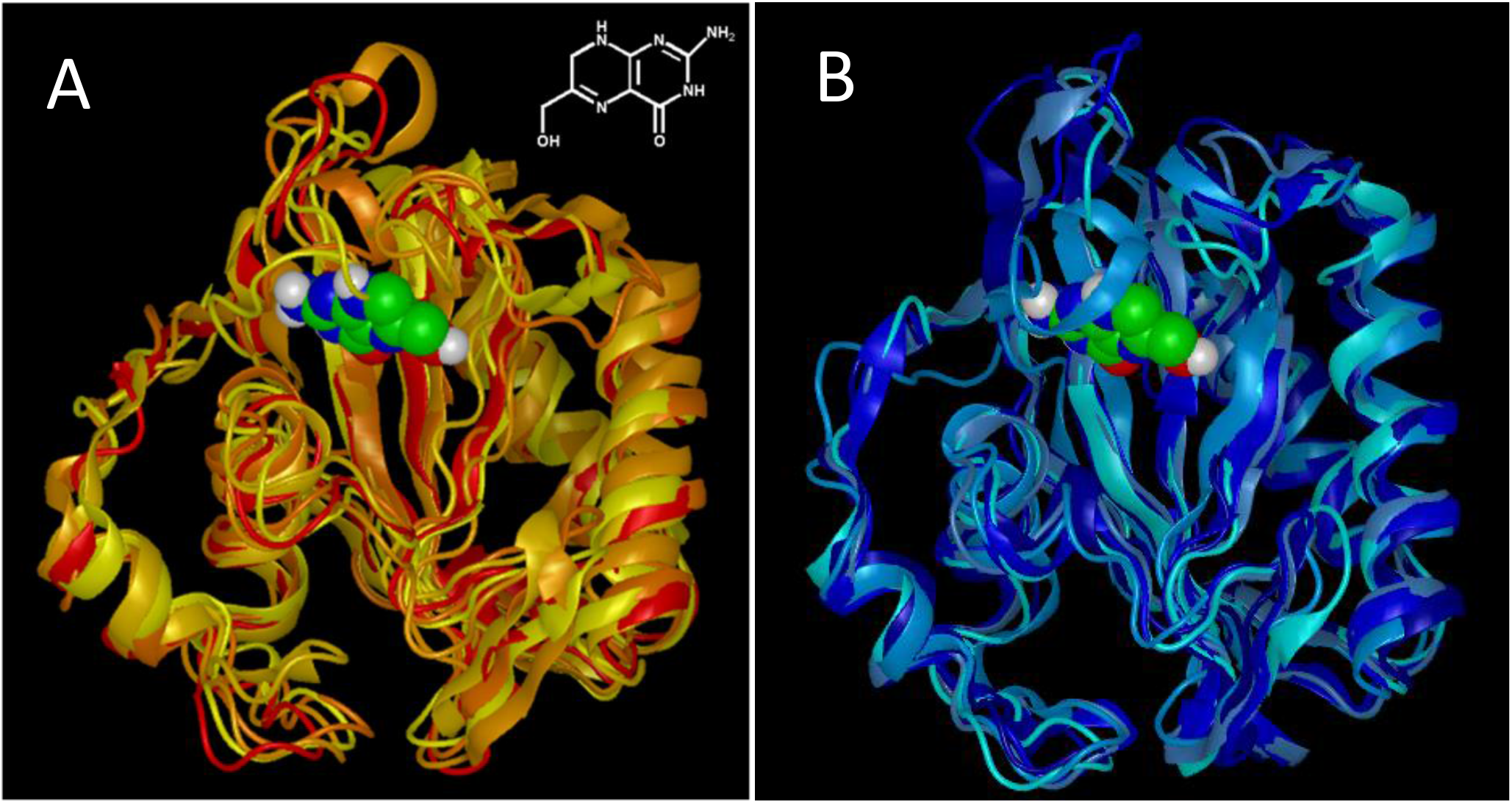
The five most populated conformational clusters from HREM molecular dynamics simulation of HPPK. Clustering with G_cluster from the GROMACS package http://www.gromacs.org; [47]) was performed using a 1.5 Å threshold on main-chain RMSD value between any pair of structures in each cluster. The most populated cluster is shown with the lightest color ribbon, with darker colors used for successively less populated clusters. Molecular dynamics simulations were performed in the absence of the dihydropterin ligand (shown here for reference, in atom-colored CPK representation: carbon - green, nitrogen - blue, hydrogen - white, oxygen - red; chemical structure is shown at upper right in panel A). **(A)***E coli* HPPK main-chain conformers, starting from the closed, deligated crystal structure (PDB entry code 1Q0N). **(B)** *Y pestis* HPPK conformers, starting from the deligated form of PDB crystal structure 2QX0.

### Defining occupied/available volumes surrounding the pterin binding site in the dynamics trajectories

A grid with 0.5 Å spacing was overlaid on the HREM MD and ROCK dynamics trajectories, which were superimposed into the same reference frame via the rigid beta sheet scaffold of HPPK (see Experimental Methods). In each of the 2000 MD and 400 ROCK snapshots, each vertex in the grid was labeled according to whether or not a 1.4 Å probe sphere (representing a potential ligand atom) could occupy that position without interpenetrating the protein (Figure 4(A)). Upon doing this for the entire MD or ROCK trajectory plus the starting closed crystallographic conformation, each vertex in the grid was labeled according to the percentage of the trajectory in which the probe could occupy that position (Figure 4(B)). The results can also be visualized as a series of molecular surface shells contoured by protein atom occupancy (Figure 5).

**Figure 4.**
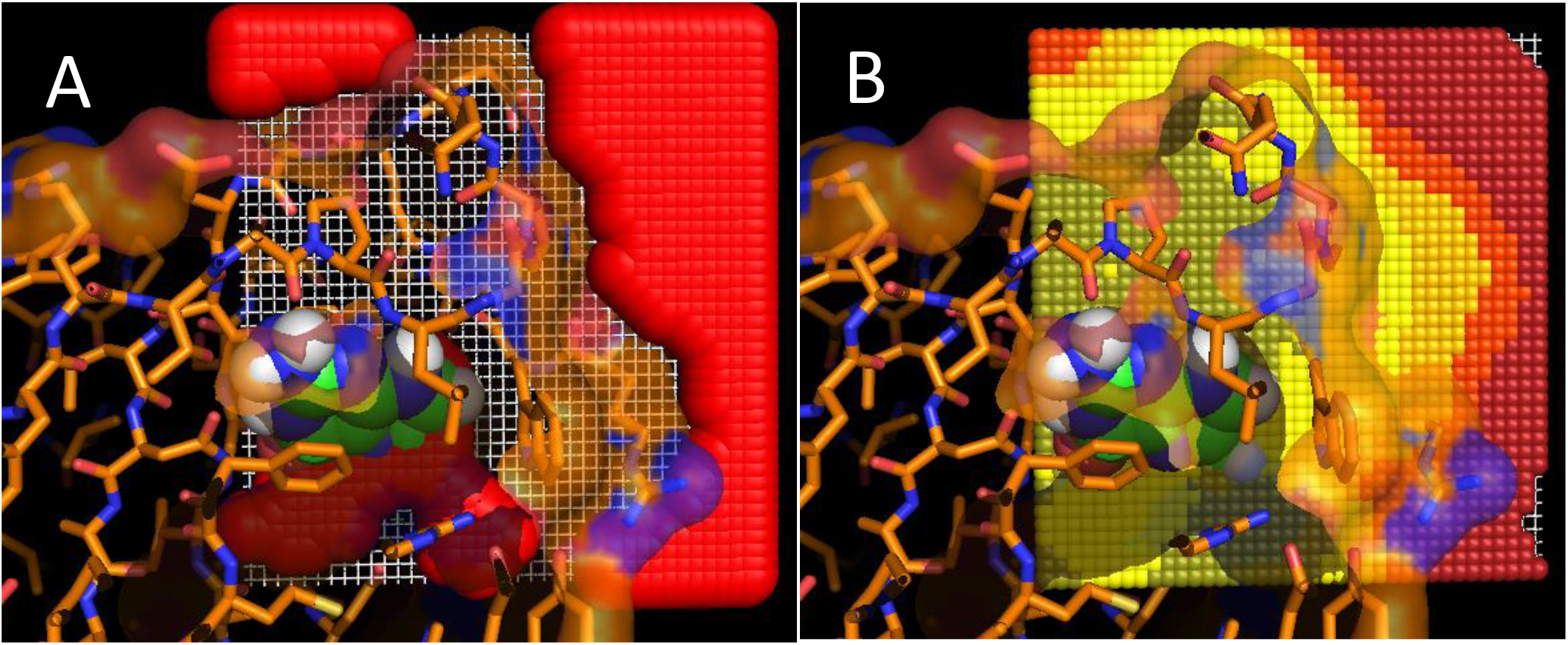
Defining occupied/available volumes surrounding the pterin binding site in the dynamics trajectories. For reference, a section of the *E coli* closed crystal structure (PDB entry 1Q0N) is shown in tube representation, with carbon atoms colored orange, and its transparent solvent accessible molecular surface is colored by atom type. Dihydropterin appears in CPK representation, with carbon atoms in green. **(A)** Probe spheres with a radius of 1.4 Å were placed on every point of a 0.5 Å grid superimposed on the dynamics trajectory. For each snapshot in the trajectory, probe spheres exhibiting van der Waals collisions with any protein atoms were removed. The resulting set of probe spheres, shown for one layer in the grid as overlapping bright red spheres surrounding the right edge of the protein, represent positions available for occupancy by ligand atoms. This analysis was performed for all snapshots in a dynamics trajectory and for all layers in the 3-dimensional grid surrounding the dihydropterin binding site. **(B)** Following analysis of the entire *E* coli HPPK molecular dynamics trajectory, each grid point is colored based on the percentage of the trajectory a probe was able to be placed in that position without van der Waals overlaps. Gold points indicate that probes could never placed at that grid point (0% availability), and yellow, orange, red, and dark red indicates points where probes could be placed 25%, 50%, 75%, and 100% of the time, respectively.Percentage availability is the complement of percentage occupancy. For example, a grid point that was occupied 25% of the time is considered to be 75% available. Empty grid points at the extreme right edge indicate where a probe could always be placed, and therefore where protein atoms never interacted.

**Figure 5.**
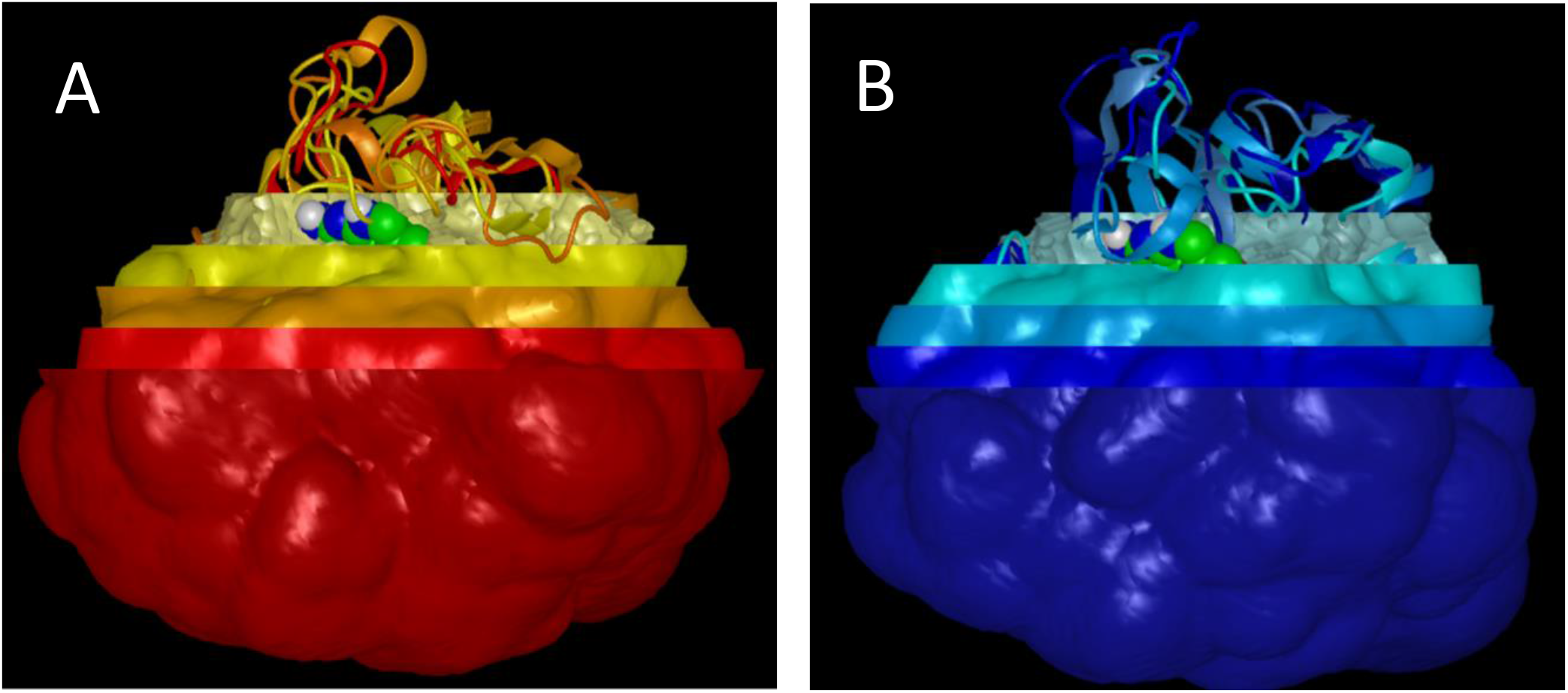
“Russian doll” representation of nested surface shells isocontoured around volumes that were 100% occupied (most central, brightest colored shell; 0% available), 75% occupied (next most central shell), then 50%, 25%, and 0% occupied (outermost shell) over the entire HREM trajectory. Conformational cluster representatives and the dihydroperin site from Figure 3 are shown for reference. **(A)** *E coli* HPPK occupied volumes in layers from gold (100% occupied) to red (0% occupied). **(B)** *Y pestis* HPPK occupied volumes from silver (100% occupied) to dark blue (0% occupied).

### Defining a unique available volume to gain conformational selectivity for the target protein

A unique volume to gain selectivity for *Y pestis* HPPK needed to be available a significant percentage of time, such that the energetic cost of opening the pocket volume was not significant, e.g., ~1 kcal/mol or less (Table 1). There are essentially three criteria to be considered when selecting a unique pocket volume neighboring the substrate site. It should be: (i) able to accommodate several ligand atoms in a pocket that maximizes ligand interactions, (ii) significantly available over time as a function of the protein’s dynamics, and (iii) unavailable or rarely available to ligand atoms in the off-target homolog. The trade-off between these factors is determined by the conformational dynamics of the two proteins being considered. Here, it was possible to select the shell of *E coli* HPPK that is entirely occupied in all the MD and ROCK conformers (Figure 5), overlay the 25% available shell of *Y pestis* HPPK, and identify a volume subtended by these surfaces and adjacent to the dihydropterin binding site that could accommodate several atoms in *Y pestis* HPPK (Figure 6). In general, one would start with the 100% occupied volume in the off-target homolog, and overlay each available-volume surface in the target protein in turn (e.g., 100% available, 75% available, 25% available, etc.) until a significantly available pocket was identified neighboring the substrate site. If the criteria could not be met with the 100% occupied surface in the off-target homolog, one would consider a slightly less-occupied shell (e.g., 90%) along with the most-available shell in the target protein that provides a unique pocket. The difference in energy of opening the pocket between the two homologs must strongly favor the target protein, according to the criteria in Table 1.

The *Y pestis* HPPK structure chosen for virtual screening was the one best complementing the surface contours of the unique and substrate-binding pockets. Based on scanning the 2000 MD snapshots, 400 ROCK snapshots, and the closed crystal structure of *Y pestis* HPPK, conformation 194 from the ROCK simulation was identified as optimal and used to derive a screening template for SLIDE. One set of key points was placed in the deepest part of the dihydropterin pocket (as described in the Experimental Methods), and the second set was defined as all template points within the unique *Y pestis* pocket. Docked molecules were required to match at least one template point in each set.

### Screening for compounds matching the substrate and unique sites

The SLIDE screening run was designed to identify molecules that: i) docked into both the dihydropterin and unique volumes, ii) had predicted affinity scores at least 2 standard deviations above the mean score (determined by screening the first 20% of the ZINC lead-like compounds), and iii) made at least three hydrogen bonds in common with the dihydropterin substrate in *Y pestis* HPPK but not in the *E coli* pterin pocket. This third criterion (step 2-4 in Figure 2) involved evaluating ligand candidates against representatives from all MD and ROCK conformational clusters for *E coli* HPPK (including the closed structure), and was designed to rule out compounds that could bind well in a different mode to the dihydropterin pocket of *E coli*.

### Top-ranking compounds identified by the SpeciFlex protocol

Of the 337 compounds meeting the above criteria, nineteen were found to make at least 4 hydrogen bonds in common with dihydropterin and at least one favorable interaction in the unique pocket of *Y pestis* HPPK. Eleven of the nineteen formed at least one hydrogen bond in the unique pocket. To illustrate the quality of some of the top-ranked molecules resulting from the SpeciFlex protocol, five representative compounds are shown (Figure 7), all of which shared at least three hydrogen bonds with dihydropterin and made at least two favorable interactions in the unique pocket. However, with an inhibitor hit rate of 15-19% in past applications of SLIDE (see section on Step 2-2), we recommend that 30 or more candidates be prioritized and assayed in the lab when possible.

Compound 1 is 4-(2-(2-hydroxy-3-(hydroxymethyl)-5-methylphenyl)-2-oxoethyl)-2,6-piperidine-dione (Figure 7(A); ZINC entry 1636542 and National Cancer Institute (NCI) entry 658411). At NCI, this compound was tested for anti-HIV activity and found inactive. A triply substituted phenyl group is placed in the unique pocket, forming two hydrogen bonds and two hydrophobic contacts. In the pterin pocket, four of the dihydropterin hydrogen bonds are mimicked by the piperidine-dione moiety. The predicted affinity for *Y pestis* HPPK is −8.5 kcal/mol.

Compound 2, 3,3’,4’,5,7-pentahydroxy-2-phenylbenzopyrylium chloride, also known as cyanidin (Figure 7(B); ZINC entry 3775158; NCI entry 407291), has a predicted affinity of −8.5 kcal/mol. It is hydrogen-bond acceptor-rich, yet substantially mimics dihydropterin sterically and places its hydrogen-bonding groups in similar positions. In the unique pocket (upper right), two hydrogen bonds are formed by a single hydroxyl group on the phenyl ring, with the other hydroxyl group interacting with solvent. This compound is toxic according to the ChEMBL database (https://www.ebi.ac.uk/chembldb/index.php/compound/inspect/CHEMBL404515), causing morphological changes in human HMC1 cells [47] at 50 μM concentration, and it inhibits human cyclooxygenase 1 activity by 39% at 5 μM concentration [48]. Thus, it remains important to check high-ranking compounds from virtual screening for reported biological activities and absorption, distribution, metabolism, excretion, and toxicology properties before further testing.

Compound 3 is 2-(3-methoxyphenyl)-3,4-dimethyl-2,6-dihydro-7H-pyrazolo[3,4-d]pyridazin-7-one (Figure 7(C); ZINC entry 1016729; Ambinter (vendor) entry AO-476/43249978). This compound also docks with a predicted affinity of −8.5 kcal/mol. The pyridazinone moiety is reasonably isosteric with dihydropterin and mimics its hydrogen bond network via positioning nitrogen atoms similarly. The hydroxymethyl group in the unique pocket makes one hydrogen bond and one hydrophobic contact with the protein. Like compound 2, much of its phenyl ring is solvent-exposed.

Compound 4, 3-(2-hydroxy-3-methoxyphenyl)-1-(2-hydroxyphenyl)-2-propen-1-one (Figure 7(D); ZINC entry 4252568; NCI entry 64614) has a predicted affinity of −8.6 kcal/mol and is the most flexible of the top-scoring molecules. A hydroxymethyl group mimics three of the deep-pocket hydrogen bonds of dihydropterin, and three hydrophobic contacts are formed in the dihydropterin pocket. The two dihydroxymethyl groups in the unique pocket form two hydrogen bonds and two hydrophobic contacts.

Finally, compound 5, which is 4-[(3,4-dimethylphenyl)acetyl]-3,4-dihydro-2(1H)-quinoxalinone (Figure 7(E); ZINC entry 11025824), has a predicted affinity of −8.6 kcal/mol. It is perhaps the most surprising candidate because its quinoxalinone ring system binds in-plane with the dihydropterin substrate, but is rotated by 90 degrees. It still binds deeply and replicates three of the dihydropterin hydrogen bonds to HPPK. The dimethylphenyl group in the unique pocket forms three hydrophobic interactions with HPPK.

**Figure 6.**
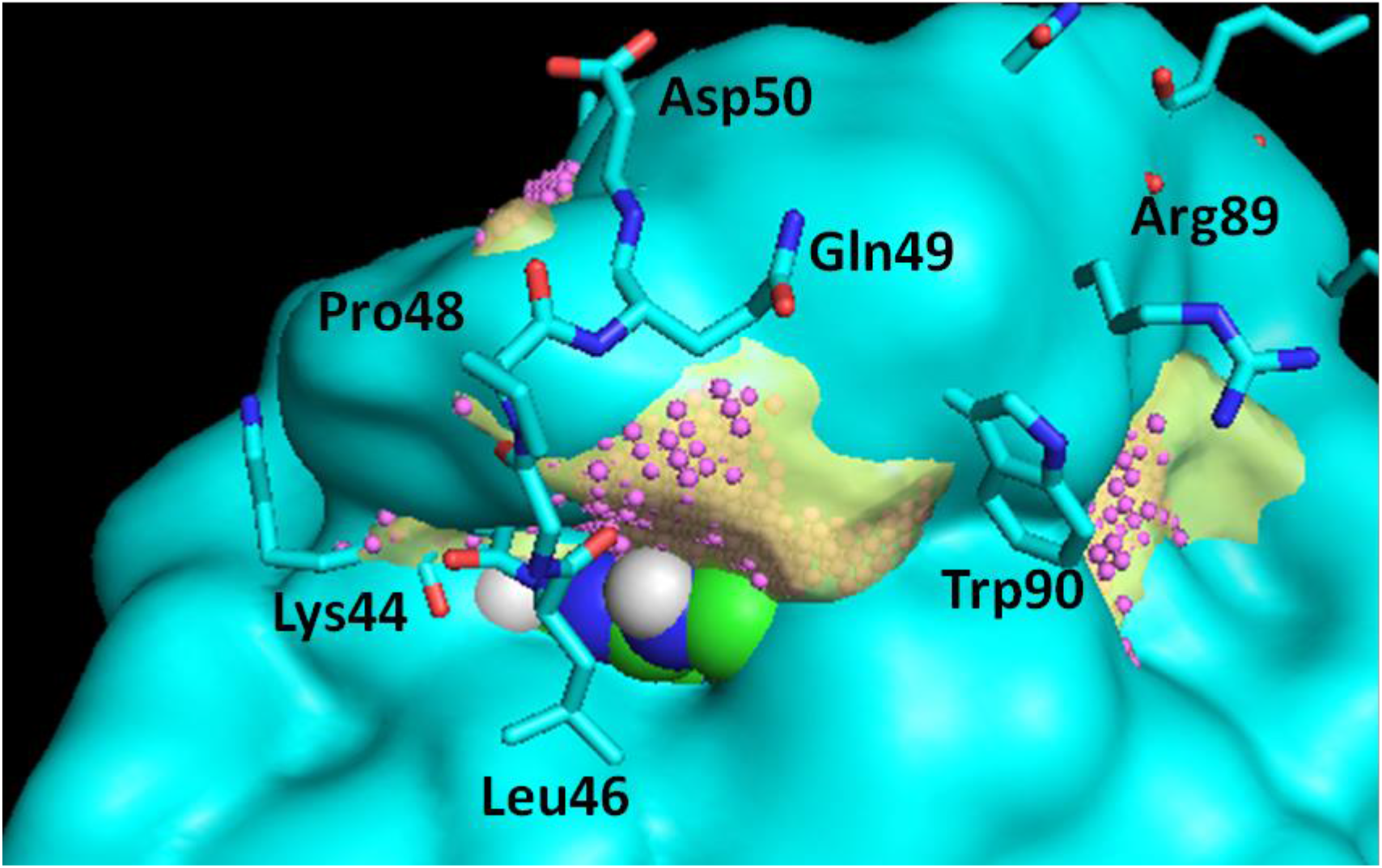
Uniquely sampled pocket adjacent to the dihydropterin pocket in *Y pestis* HPPK. The cyan surface represents the surface contour outside of which the grid points are 25% available across the *Y pestis* HREM trajectory; the interior of this contour is occupied by protein atoms 75% of the time. The transparent gold surface represents the contour containing all of the grid points where a probe was never able to be placed in the *E. coli* HPPK trajectory. The small pink spheres show all grid points in the volume between these two surfaces, representing points where a ligand atom with 1.4 Å radius could be placed within the 25% available *Y pestis* HPPK volume and within the 0% available *E coli* HPPK volume. The gold volume at center, immediately adjacent to the dihydropterin site (shown in CPK model), was selected as the unique pocket targeted for compounds that could bind to *Y pestis* HPPK while being disallowed from binding in the corresponding region of *E coli* HPPK.

**Figure 7.**
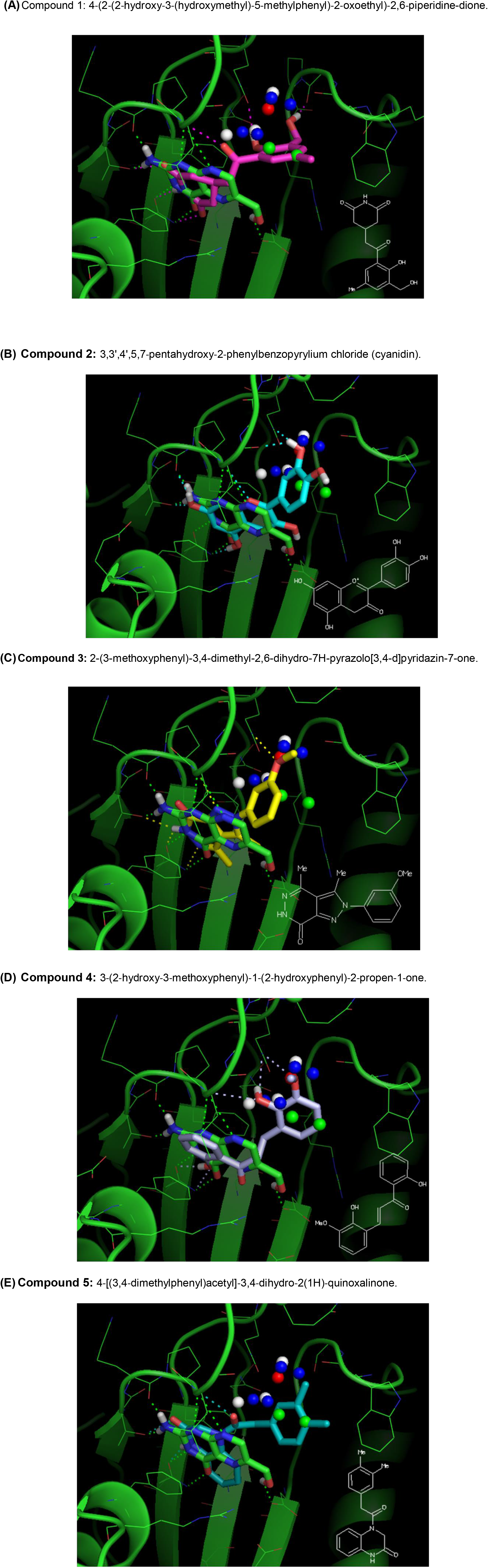
The target structure of *Y pestis* HPPK (ROCK conformer 194) is shown with each docked top-ranking molecule. The position of dihydropterin from the closed crystal structure is shown for reference (tubes with green carbon atoms). Template point spheres representing ideal hydrophobic (green), hydrogen-bond donor (blue), hydrogen-bond acceptor (red) or hydrogen bond donor/acceptor (white) interaction sites for ligand atoms are shown in the unique pocket. Hydrogen bonds and salt bridges are displayed as dashed lines and side chains are displayed in their conformations following SLIDE docking. The 2dimensional structure of each compound is shown as an inset in the lower-right corner. **(A)** Compound 1: 4-(2-(2-hydroxy-3-(hydroxymethyl)-5-methylphenyl)-2-oxoethyl)-2,6-piperidine-dione. **(B)** Compound 2: 3,3',4',5,7-pentahydroxy-2-phenylbenzopyrylium chloride (cyanidin). **(C)** Compound 3: 2-(3-methoxyphenyl)-3,4-dimethyl-2,6-dihydro-7H-pyrazolo[3,4-d]pyridazin-7-one. **(D)** Compound 4: 3-(2-hydroxy-3-methoxyphenyl)-1-(2-hydroxyphenyl)-2-propen-1-one. **(E)** Compound 5: 4-[(3,4-dimethylphenyl)acetyl]-3,4-dihydro-2(1H)quinoxalinone.Figure 7,.

## Conclusions

The SpeciFlex protocol described here consists of one set of steps for identifying a unique pocket in the target protein relative to its homolog based on differential protein dynamics, and a second set of steps for using this information to screen for inhibitor candidates that complement the unique pocket and a neighboring substrate site. Results show that the resulting high-scoring, low-molecular weight (≤ 300D) candidates interact well with both the unique and substrate pockets. Cross-docking was used to filter out any compounds that could interact similarly well with the off-target homolog. Thus, a smooth workflow now exists for mining and visualizing conformational population differences between homologs, and applying this information to virtual screening. The top-scoring candidates are chemically diverse while sharing some features: they mimic dihydropterin interactions via an isosteric or alternative ring system, which is connected by a short flexible linker to a second cyclic group binding in the unique pocket of *Y pestis* HPPK.

## Executive Summary

- Molecular dynamics analysis allows identification of conformational sampling differences between homologous proteins that otherwise have very similar binding sites in the closed conformation
- Considering the difference in population of a targeted conformation between two homologs can provide guidance on whether conformational selectivity is a feasible way of enhancing ligand specificity.
- The SpeciFlex protocol allows conformational sampling differences to be identified and exploited during virtual screening. Compounds are identified that can bind well in a substrate pocket and an adjacent selectivity pocket. The compounds are also filtered to remove any that might bind similarly well to the corresponding site in the homolog.
- Steps in the procedure can be substituted by similar methods based upon user preference. For instance e.g., HREM molecular dynamics using the GROMOS force field could be replaced by CHARMM or AMBER molecular dynamics; SLIDE docking could be substituted by another method suitable for large-scale virtual screening, such as GLIDE; proprietary databases could be used in place of ZINC; and different scoring protocols could be used to rank the candidates. SpeciFlex is essentially a modular protocol in which the emphasis is placed on how to incorporate conformational population differences as a basis for selectivity in virtual screening.
- For the target enzyme HPPK, SpeciFlex filtered more than 800,000 lead-like compounds down to a set of 19 compounds that met the criteria of having high predicted binding affinity and good complementarity with the substrate and unique pocket sites, while exhibiting considerable diversity in chemical structures.

## Footnotes

1 The SpefiFlex software is available from GitHub at https://github.com/psa-lab/SpeciFlex

## Acknowledgements

We thank Robert Cukier and Li Su for providing HREM molecular dynamics trajectories for *Y pestis* and *E coli* HPPK, and for suggesting their cavity search approach as a starting point for analyzing occupancy in molecular dynamics trajectories. We appreciate Honggao Yan for engaging us in inhibitor discovery for HPPK and the opportunity to work together on experimental validation of the ligand candidates presented here. Maria Zavodszky, Sandeep Namilikonda, and John Johnston made valuable extensions to the ROCK software. OpenEye Scientific Software (Santa Fe, NM) has generously provided its Vida, QuaCPac, Omega, Syzbki, and OEChem software for use in this project. The Pfizer Strategic Alliance program, the Michigan State University Quantitative Biology Initiative, and the MSU College of Natural Science provided funding for this research.

